# Saccharibacteria harness light energy using Type-1 rhodopsins that may rely on retinal sourced from microbial hosts

**DOI:** 10.1101/2022.02.13.480300

**Authors:** Alexander L. Jaffe, Masae Konno, Yuma Kawasaki, Chihiro Kataoka, Oded Béjà, Hideki Kandori, Keiichi Inoue, Jillian F. Banfield

**Affiliations:** Department of Plant and Microbial Biology, University of California, Berkeley, CA; The Institute for Solid State Physics, The University of Tokyo, Kashiwa, Chiba, Japan; PRESTO, Japan Science and Technology Agency, Kawaguchi, Saitama, Japan; Department of Life Science and Applied Chemistry, Nagoya Institute of Technology, Showa-ku, Nagoya, Aichi, Japan; Faculty of Biology, Technion-Israel Institute of Technology, Haifa, Israel; OptoBioTechnology Research Center, Nagoya Institute of Technology, Showa-ku, Nagoya, Aichi, Japan; Innovative Genomics Institute, University of California, Berkeley, CA; Department of Earth and Planetary Science, University of California, Berkeley, CA; Department of Environmental Science, Policy, and Management, University of California, Berkeley, CA

## Abstract

Microbial rhodopsins are a family of photoreceptive membrane proteins with a wide distribution across the Tree of Life. Within the Candidate Phyla Radiation (CPR), a diverse group of putatively episymbiotic bacteria, the genetic potential to produce rhodopsins appears to be confined to a small clade of organisms from sunlit environments. Here, we characterize the metabolic context and biophysical features of Saccharibacteria Type-1 rhodopsin sequences derived from metagenomic surveys and show that these proteins function as outward proton pumps. This provides one of the only known mechanisms by which CPR can generate a proton gradient for ATP synthesis. Intriguingly, Saccharibacteria do not encode the genetic machinery to produce all-*trans*-retinal, the chromophore essential for rhodopsin function, but their rhodopsins are able to rapidly uptake this cofactor when provided in experimental assays. We found consistent evidence for the capacity to produce retinal from β-carotene in organisms co-occurring with Saccharibacteria, and this genetic potential was dominated by Actinobacteria, which are known hosts of Saccharibacteria in other habitats. If Actinobacteria serve as hosts for Saccharibacteria in freshwater environments, exchange of retinal for use by rhodopsin may be a feature of their associations.

## Introduction

The sun is the dominant source of energy on Earth, and many organisms have evolved ways to use light. Only recently was it suggested that putatively symbiotic Saccharibacteria (formerly TM7) of the Candidate Phyla Radiation (CPR) may be able to use rhodopsins for proton translocation and thus energy generation [1, 2]. However, experimental evidence supporting this function was lacking. Here, we biophysically characterize these rhodopsins and explore their relevance for the metabolism of Saccharibacteria. We also consider how rhodopsins may play a role in the interactions between Saccharibacteria and their microbial hosts in sunlit environmental microbiomes.

## Results

Phylogenetic placement of Saccharibacteria rhodopsins (SacRs) shows that these sequences form a sibling clade to characterized light-driven inward and outward H^+^ pumps (Fig. 1a). We selected three phylogenetically diverse SacRs from freshwater lakes and two related, previously uncharacterized sequences from Gammaproteobacteria (*Kushneria aurantia* and *Halomonas* sp.) for synthesis and functional characterization. All sequences have Asp–Thr–Ser (DTS) residues at the positions of D85–T96–D96 of bacteriorhodopsin (BR) in the third transmembrane helix (Fig. S1). These residues are known as the triplet DTD motif and represent key residues for proton pumping function in BR [3].

**Figure 1.**
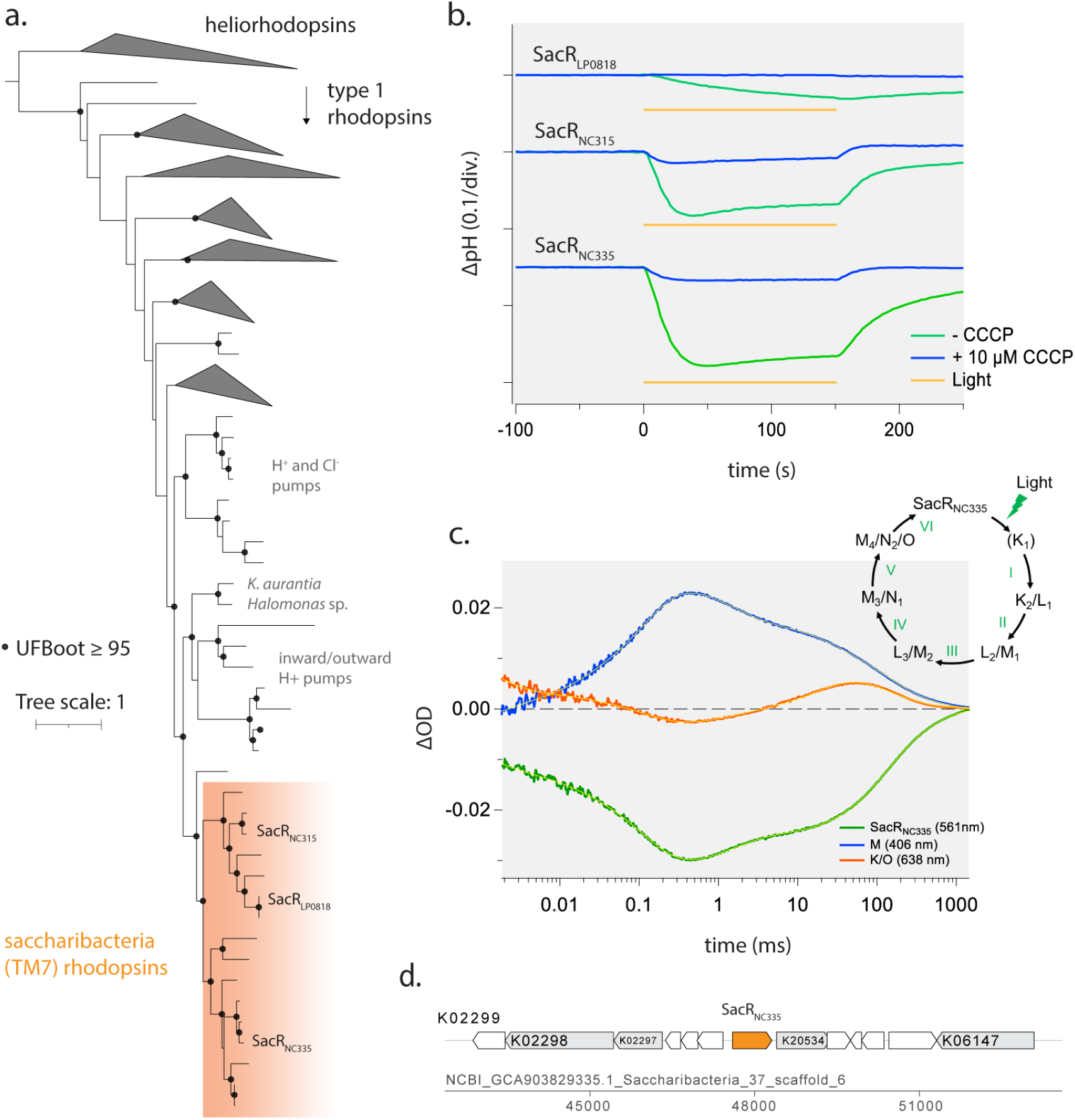
Characteristics of Saccharibacteria rhodopsins (SacRs). **a**) Rhodopsin gene tree indicating that SacRs form a broad clade of proton pumps. **b**) The ion-pumping activity of SacRs. Blue and green lines indicate the pH change with and without 10 μM CCCP, respectively. Yellow bars indicate the period of light illumination. **c**) Time evolution of transient absorption changes of SacR_NC335_ in 100 mM NaCl, 20 mM HEPES–NaOH, pH 7.0, and POPE/POPG (molar ratio 3:1) vesicles with a lipid to protein molar ratio = 50. Time evolution at 406 nm (blue, representing the M accumulation), 561 nm (green, representing the bleaching of the initial state and the L accumulation) and 638 nm (red, representing the K and O accumulation). Yellow lines indicate fitting curves by a multi-exponential function. **Inset:** The photocycle of SacR_NC335_ based on the fitting in **c)** and a kinetic model assuming a sequential photocycle. The lifetime (*τ*) of each intermediate is indicated by numbers as follow (mean ± S.D., fraction of the intermediate decayed with each lifetime in its double exponential decay is indicated in parentheses): I: *τ* = 1.7 ± 0.3 ms (42%), *τ* = 13 ± 1.8 ms (58%), II: *τ* = 118 ± 2 ms, III: *τ =* 1.6 ± 0.1 ms, IV: *τ* = 23.5 ± 1.0 ms, V: *τ* = 98.4 ± 6.4 ms (56%), *τ* = 384 ± 18 ms (44%). **d)** Genomic context of SacR_NC335_. Neighboring genes with above-threshold KEGG annotations are indicated in gray with the highest-scoring HMM model. Genes without KEGG annotations are indicated in white.

Proton transport assays for the SacRs and Gammaproteobacteria proteins expressed in *E. coli* showed marked decrease of external pH upon light illumination (Fig. 1b, Fig. S2), indicating that these proteins are light-driven outward H^+^ pumps. The pH decrease was almost eliminated after adding the protonophore carbonyl cyanide m-chlorophenyl hydrazone (CCCP), which dissipates the H^+^ gradient, confirming that it was indeed formed upon illumination (Fig. 1b, Fig. S2). We also characterized the absorption spectra and the photocycle of the SacRs, showing that the three rhodopsins have an absorption peak around 550 nm (Fig. S3). The photocycle of the SacRs, determined by measuring the transient absorption change after nanosecond laser pulse illumination (Fig. 1c, Fig. S4), displays a blue-shifted M intermediate that represents the deprotonated state of the retinal chromophore. This has been observed for other H^+^ pumping rhodopsins [4, 5] and indicates that the proton bound to retinal is translocated during pumping.

Given that SacRs function as outward proton pumps, we searched Saccharibacteria genomes for the F_1_F_o_ ATP synthase that would be required to harness the generated proton motive force for ATP synthesis. HMM searches showed that all genomes encoded the complete ATP synthase gene cluster and, furthermore, had c subunits with motifs consistent with H^+^ binding, instead of Na^+^ binding (Table S1, Fig. S5). Together, our experimental and genomic analyses strongly suggest that some Saccharibacteria utilize rhodopsins for auxiliary energy generation in addition to their core fermentative capacities.

Retinal is the rhodopsin chromophore that enables function of the complex upon illumination [6]. Intriguingly, we found no evidence for the presence of β-carotene 15,15’-dioxygenase (*blh*), which produces all-*trans*-retinal (ATR) from β-carotene, in Saccharibacteria genomes with rhodopsin. This absence was likely not due to genome incompleteness, as rhodopsin genomic loci were well-sampled and neither *blh* nor any conserved hypothetical proteins were present in these regions, where *blh* is often found [7] (Fig. 1d, Fig. S6, Table S2). As SacRs do contain the conserved lysine for retinal binding [1], we instead hypothesized that Saccharibacteria may uptake retinal from the environment, as has been previously observed for some organisms with rhodopsin that also lack *blh* [8, 9].

We next tested the ability of SacR proteins to bind ATR from an external source by performing a retinal reconstitution assay. In contrast to the proton transport assays, where rhodopsin was expressed in the presence of ATR, here ATR was dissociated from the purified complex and the visible absorbance of rhodopsin was measured upon re-addition of ATR [10]. Both *Gloeobacter* rhodopsin (GR), a typical Type-1 outward H^+^ pump, and SacRs showed an increase in absorption in the visible region with time after the addition of ATR (Fig. 2a, Fig. S7). For all SacRs, the binding of ATR by their apoprotein was saturated within 30 seconds after retinal addition (Fig. 2b), indicating that SacR is able to be efficiently functionalized using externally derived ATR. The observed reconstitution rate is substantially faster than that of GR (> 20 minutes) and comparable to that of heliorhodopsin, which is used by other microorganisms also lacking a retinal synthetic pathway and rapidly binds ATR through a small opening in the apoprotein [9]. In the structure of SacR_NC335_ modeled by Alphafold2 [11, 12], a similar hole is visible in the protein moiety near the retinal (Fig. S8). Hence, SacRs might also bind retinal through this hole in a similar manner to *T*aHeR (heliorhodopsin).

**Figure 2.**
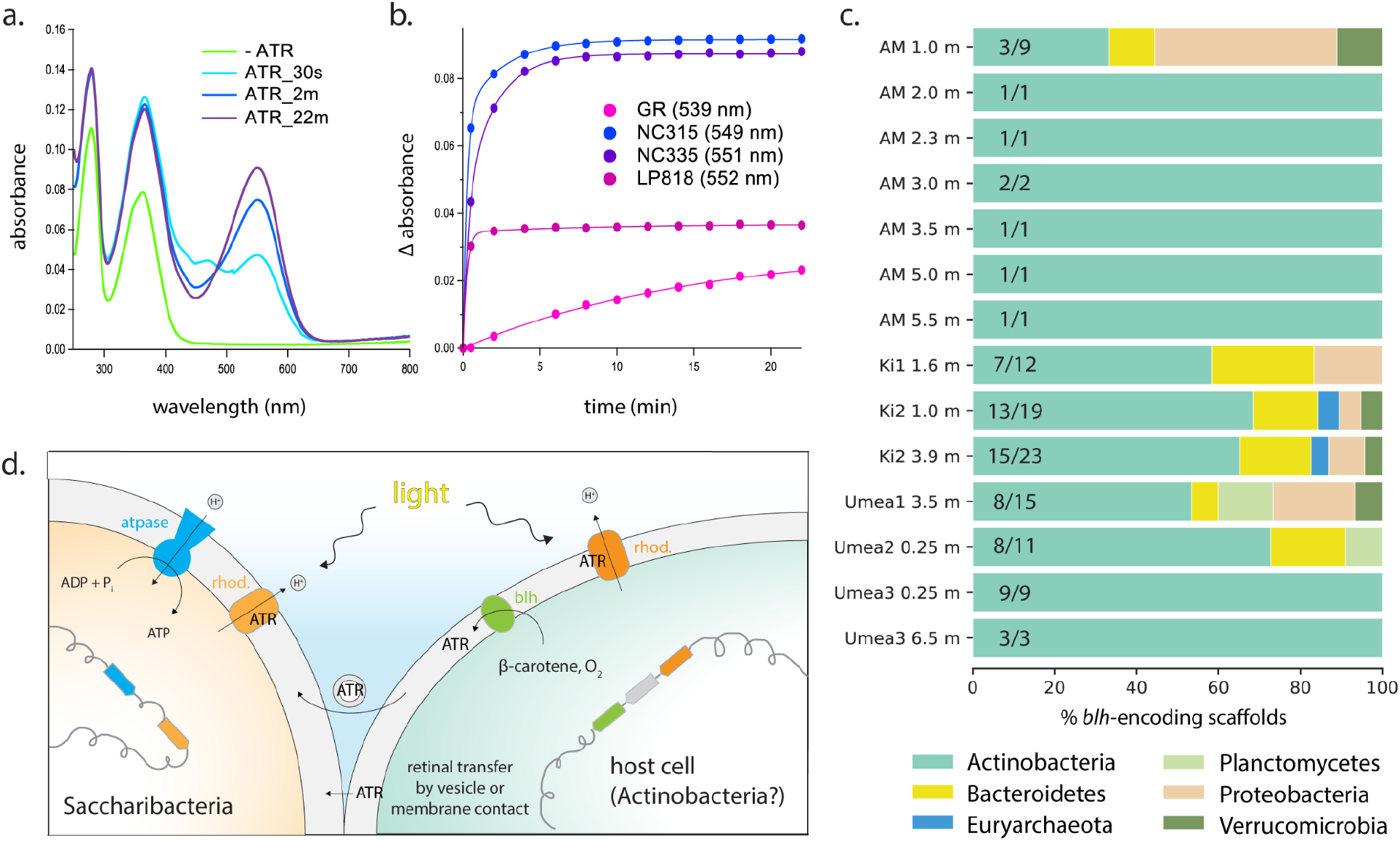
Binding of retinal by Saccharibacteria rhodopsins and context for biosynthesis. **a**)UV-visible absorption spectra showing the regeneration of retinal binding to SacR_NC335_ and GR in 20 mM HEPES–NaOH, pH 7.0, 100 mM NaCl and 0.05% n-dodecyl-β-D-maltoside (DDM). In SacR_NC335_, a peak around 470 nm was transiently observed in the spectrum 30 seconds after the addition of ATR, suggesting that an intermediate species appears during the retinal incorporation process that involves formation of the Schiff base linkage. **b**) Time evolution of visible absorption representing retinal binding to apo-protein. Numbers in parentheses in the legend indicate the absorption maxima of each rhodopsin. **c**) Genetic potential for β-carotene 15,15’-dioxygenase (*blh*) production in freshwater lake metagenomes where SacRs are found. Fractions indicate the number of *blh*-encoding scaffolds taxonomically affiliated with Actinobacteria. **d**) Conceptual diagram illustrating potential retinal exchange between Saccharibacteria and host cells. Abbreviations: ATR, all-*trans-*retinal; GR, *Gloeobacter* rhodopsin; AM, Alinen Mustajärvi; Ki, Kiruna; rhod., rhodopsin.

Saccharibacteria with rhodopsin must obtain retinal from other organisms. To evaluate possible sources of ATR, we investigated the genetic potential for retinal biosynthesis in 15 subarctic and boreal lakes [13] where Saccharibacteria with rhodopsin were present (Fig. S9). *Blh*-encoding scaffolds were found in 14 of the 15 metagenomes profiled (∼93%), and, in nearly all cases these scaffolds derived from Actinobacteria (Fig. 2c, Table S3). This is intriguing because Actinobacteria are known to be hosts of Saccharibacteria in the human microbiome [14, 15] and potentially more generally [1, 16]. BLAST searches against genome bins from the same samples indicated that these Actinobacteria were members of the order Nanopelagicales (Table S3), and often encoded a rhodopsin (phylogenetically distinct from SacRs) in close genomic proximity to *blh* genes (Table S4). HMM searches revealed that these genomes also harbor homologs of the *crtI, crtE, crtB*, and *crtY* genes necessary for β-carotene production [17].

## Discussion

Here, we add to growing evidence that DTS-motif rhodopsins can function as outward H^+^ pumps [18] and report that Saccharibacteria use them to establish a proton gradient used for energy generation, given a source of ATR and light. This is one of the very few known ways that any CPR organism can pump protons across the membrane. However, the source of the ATR enabling function of Saccharibacteria rhodopsins is unclear. While there is precedent for external supply of ATR to functional rhodopsins in other bacteria [8, 9], the mechanism by which this hydrophobic compound is transferred to the membrane of such bacteria is also unknown.

Experimental co-cultures of Saccharibacteria with Actinobacteria from multiple microbiome types [15, 16] suggest that a host bacterium for the Saccharibacteria studied here may be the source of ATR. We infer that these hosts are co-occurring Nanopelagicales Actinobacteria that dominate retinal production in microbial communities containing Saccharibacteria with rhodopsin. Retinal produced by these Actinobacteria from β-carotene may be transferred to Saccharibacteria either by membrane contact, a common feature in imaged CPR-host interactions [14, 19], or possibly via extracellular vesicles (Fig. 2d). ATR produced by Actinobacteria is required for their own rhodopsins [8] (Fig. 2d), but it is conceivable that they (if they indeed are the hosts) produce ATR in excess to deliberately supply Saccharibacteria symbionts, possibly to ensure interdependence. Alternatively, Saccharibacteria scavenge ATR. Regardless of the source organism and ATR transfer mechanism, our analyses suggest a new aspect of Saccharibacteria lifestyles, where they employ rhodopsins and externally derived retinal to produce energy via phototrophy.

*For Materials and Methods, see Supplementary Methods*.

### Data and software availability

All accession information for the genomes and metagenomic samples analyzed in this study are listed in the Supplementary Tables. Additional files (including the masked rhodopsin alignment and maximum likelihood tree), supplementary tables, and custom code for the described analyses are also available on Zenodo (doi.org/10.5281/zenodo.6038621).

## Acknowledgments

We thank Marie C. Schoelmerich for helpful discussions and guidance on metabolic analyses. We thank Anne-Catherine Lehours and the Laboratoire Microorganismes: Génome Environnement at Université Clermont Auvergne for their assistance in generating the newly-reported Lac Pavin sequence. Funding was provided by Moore Foundation Grant 71785 to J.F.B. This work was also supported by MEXT KAKENHI, Grant-in-Aid for Transformative Research Areas (B) “Low-energy manipulation” (Grant Number: JP20H05758 to K.I), JSPS KAKENHI, Grants-in-Aid (Grant Numbers: JP21H01875, JP20K21383 to K.I.). We also thank the Innovative Genomics Institute at UC Berkeley. O.B. holds the Louis and Lyra Richmond Chair in Life Sciences.

## Author contributions

A.L.J., O.B., K.I., and J.F.B. designed the project. A.L.J. and K.I. performed bioinformatic and phylogenetic analyses. M.K., Y.K., C.K., and H.K. performed biophysical assays. A.L.J., J.F.B., K.I., and M.K. wrote the manuscript. All authors made comments on the manuscript.

## SUPPLEMENTARY INFORMATION

## Supplementary Methods

### Phylogenetic tree building

Rhodopsin protein sequences from Jaffe et al. 2021 [1] were combined with additional reference sequences from public databases as well as a set of newly-reported sequences from freshwater Saccharibacteria for phylogenetic analysis [2]. Also included was a representative rhodopsin sequence from a freshwater Saccharibacteria genome derived from a metagenomic survey of the meromictic Lac Pavin. Identical sequences were removed using usearch (*-cluster_fast -id 1*) [20] and aligned using ClustalW [21]. Alignment of the 1st, 2nd, and 5th transmembrane helices were corrected manually. Finally, a maximum likelihood tree was inferred using IQTree (*-m TEST -bb 1000*) [22]. The corrected alignment and maximum likelihood tree are available on Zenodo (see **Data and Software availability** statement).

### Sequence synthesis and expression

Three phylogenetically diverse Saccharibacteria rhodopsin sequences and two Gammaproteobacteria sequences were selected for experimental characterization. To ensure the accuracy of protein predictions, metagenomic reads for each sample were mapped back to assembled Saccharibacteria contigs with rhodopsin using bowtie2 (default parameters) [23]. Read mapping for each contig was examined manually using Geneious.

Genes encoding each rhodopsin were artificially synthesized with codon optimization for an *E. coli* expression (Genscript, Piscataway, NJ, USA) and cloned into *Nde*I-*Xho*I site of pET21a(+) vector (Novagen, Merck KGaA, Germany). The constructed plasmids were transformed into *E. coli* C43(DE3) strain (Lucigen, WI, USA). The protein expression was induced by 0.1 mM isopropyl-β-D-thiogalactopyranoside (IPTG) in the presence of 10 μM ATR (Toronto Research Chemicals, Canada) for 4 h at 37 °C. The expressed proteins had a 6 × His-tag on the C-terminus.

### Proton transport, absorption spectra, and photocycle of Saccharibacteria rhodopsins

Light-driven proton transport was measured according to previously reported protocol [24] . Briefly, the number of *E. coli* cells expressing each rhodopsin was estimated by their optical density of the culture at 660 nm (OD_660_). In the following measurements, the equivalent of 15 OD_660_•mL (OD_660_ × total volume (ml) of the cell suspension) cell amount was used. The cells were collected by centrifuge (4,800 × *g*, 2 min, 20 °C), washed once and equilibrated three times with unbuffered 100 mM NaCl solution. The equilibrated cells were resuspended by unbuffered 100 mM NaCl solution and adjusted at OD_660_ = 2. The cell suspension was placed in the dark at 20 °C and illuminated at *λ* > 500 nm using a 300 W xenon light source (MAX-303, Asahi Spectra, Japan) through a long pass filter (Y-52; AGC Techno Glass, Japan) and a heat-absorbing filter (HAF-50S-50H; SIGMAKOKI, Japan). The light-induced changes in Ph were measured using a pH electrode (9618S-10D; HORIBA, Japan). Measurements were repeated under the same condition after the addition of 10 μM CCCP.

Proteins were purified using a 5 mL Co^2+^-NTA column (HiTrap TALON crude; Cytiva,Tokyo, Japan) on an ÄKTA start protein purification system (Cytiva, MA). The rhodopsin-expressing cells were harvested and resuspended in a buffer containing 50 mM Tris–HCl (pH 8.0) and 5 mM MgCl_2_. The harvested cells were disrupted by sonication (Ultrasonic Homogenizer VP-300N, TAITEC, Japan). The membrane fraction was collected by ultracentrifugation (CP80NX, Eppendorf Himac Technologies, Japan) at 142,000 × *g* for 1 h. The proteins were solubilized in a buffer containing 50 mM Tris–HCl (pH 7.5), 300 mM NaCl, 5 mM Imidazole and 3 % n-Dodecyl-β-D-maltopyranoside (DDM). Solubilized proteins were separated from insoluble fractions by ultracentrifugation at 142,000 × g for 1 h. After loading the solubilized proteins on Co^2+^-NTA column, the column was washed with a buffer containing 50 mM Tris–HCl (pH 7.5), 300 mM NaCl, 15 mM Imidazole and 0.1 % DDM. The His-tagged proteins were eluted with a buffer containing 50 mM Tris–HCl (pH 7.5), 300 mM NaCl, 500 mM Imidazole and 0.1 % DDM. The eluted proteins were dialyzed using buffer 20 mM HEPES–NaOH (pH 7.0), 100 mM NaCl, 0.05 % DDM to remove imidazole. Absorption spectra were recorded with a UV–vis spectrometer (V-750, JASCO, Japan).

For the transient absorption measurement by laser flash photolysis method, purified SacRs were reconstituted into a mixture of 1-palmitoyl-2-oleoyl-phosphatidyl-ethanolamine (POPE, Avanti Polar Lipids, AL) and 1-palmitoyl-2-oleoyl-sn-glycero-3-phosphoglycerol (POPG, sodium salt, Avanti Polar Lipids, AL) (molar ratio = 3:1), with a protein to lipid molar ratio of 1:50, in buffer containing 20 mM HEPES–NaOH, 100 mM NaCl, pH 7.0. The sample was illuminated with a beam of the second harmonics of a nanosecond-pulsed Nd-YAG laser (*λ* = 532 nm, 1.4–0.5 Hz, INDI40, Spectra-Physics, CA) with a pulse energy of 4.5 mJ/cm^2^/pulse. The transient absorption spectra were obtained by monitoring the intensity change of white-light from a Xe-arc lamp (L9289-01, Hamamatsu Photonics, Japan) passed through the sample, with an ICCD linear array detector (C8808-01, Hamamatsu Photonics, Japan). To increase the signal-to-noise (S/N) ratio, 10–30 identical spectra were averaged and a singular value decomposition analysis was applied. To measure the time-evolution of transient absorption change at specific wavelengths with better time resolution, the light of Xe-arc lamp (L9289-01, Hamamatsu Photonics, Japan) was monochromated by monochromators (S-10, SOMA OPTICS, Japan) and the change in the intensity after the photo-excitation was monitored with a photomultiplier tube (R10699, Hamamatsu Photonics, Japan) equipped with a notch filter (532 nm, bandwidth = 17 nm, Semrock, NY) to remove the scattered pump pulse. To increase S/N ratio, 50–100 signals were averaged.

### Analysis of metabolism and genomic context of Saccharibacteria rhodopsins

Proteins were predicted for all rhodopsin-encoding Saccharibacteria bins using Prodigal [25] (single mode) and subjected to annotation with KofamScan [26]. HMM hits were initially filtered to those with an e-value ≥ 1×10^−6^ and the highest-scoring hit for each open reading frame was selected. These results were used to identify candidate gene clusters encoding the F_1_F_o_ ATP synthase in each genome. The presence of all subunits was verified manually using a combination of KEGG annotations and additional BLAST searches. C subunit protein sequences of these gene clusters were aligned with reference sequences and manually examined for the Q…ES/T motif indicative of Na^+^ binding [27]. Finally, genomic context of the three experimentally characterized Saccharibacteria rhodopsins was visualized using gggenes (https://github.com/wilkox/gggenes). KoFamScan annotations were secondarily filtered to those with e-value ≥ 1×10^−20^ and displayed for those open reading frames within 5 kilobases upstream or downstream of rhodopsin sequences.

### Community metabolic potential for retinal biosynthesis

To examine the composition and metabolic potential of communities with rhodopsin-encoding Saccharibacteria, we used a subset of Saccharibacteria genomes reconstructed from a prior metagenomic survey of boreal/subarctic lakes [13]. We identified the sample (or samples, in the case of co-assemblies) from which each genome was constructed and mapped quality-filtered metagenomic reads from these samples back to the corresponding genome using bowtie2 (default parameters). We next used inStrain [28] to compute per-gene read coverage and breadth (fraction of gene covered) for each genome-sample pair. Samples where any Saccharibacteria rhodopsin gene attained 80% coverage breadth (or better) were retained for downstream analyses (Fig. S9).

First, quality-filtered metagenomic reads from each sample were assembled using MEGAHIT [29] (*--min-contig-len 1000*) and all proteins were predicted using Prodigal (meta mode). Predicted metagenomic proteins were searched for β-carotene 15,15’-dioxygenase (*blh*, K21817) using hmmsearch. Proteins with high-scoring hits to the K21817 HMM (at or above the model-specific score threshold) were subjected to secondary annotation with KoFamScan to verify that there were no higher-scoring HMM models.

Next, we determined taxonomic affiliation for metagenomic contigs with *blh* genes by comparing all predicted proteins on these contigs against UniRef100 using a custom DIAMOND database (diamond blastp) [30]. Hits were filtered to those with ≥ 70% coverage of the query sequence and an e-value ≥ 1×10^−5^. We retrieved taxonomic affiliation for above-threshold hits and computed the percentage of genes on each scaffold with highest similarity to various bacterial phyla. Two scaffolds with ambiguous taxonomic affiliation were curated manually using information from UniRef and BLAST searches; all other scaffolds were assigned to the phylum-level lineage with the highest percentage of hits across annotated genes.

Finally, we attempted to match *blh*-encoding scaffolds from the Actinobacteria with binned genome sequences from the same set of metagenomic samples. Moderate to high-quality Actinobacteria MAGs from [13] were downloaded and used to construct a custom BLAST database against which *blh*-encoding scaffolds from above were compared. Local alignments were filtered to those with >90% identity and >50% coverage of the query scaffolds. *Blh*-encoding scaffolds with above-threshold genome alignments were additionally assigned bin-level taxonomy determined by the original study [13]. Actinobacteria bins were then searched for *blh*, rhodopsin, and genes involved in β-carotene synthesis using KoFamScan annotations of predicted proteins as above, except employing an e-value threshold of 1×10^−5^ to capture more distant/shorter homologs of lycopene beta-cyclase (K22502).

### Retinal reconstitution assays

SacRs and GR apo-proteins were prepared by incubating purified proteins with 500 mM hydroxylamine (HA) and then illuminating at λ > 500 nm from the output of a 300 W xenon light source (MAX-303, Asahi Spectra, Japan) through a long pass filter (Y-52; AGC Techno Glass, Japan) and a heat-absorbing filter (HAF-50S-50H; SIGMAKOKI, Japan) until the proteins were completely bleached. Absorption changes representing the bleaching of rhodopsins by hydroxylamine were monitored using a UV-vis spectrometer (V-750, JASCO, Japan).

Apo-protein samples were washed five times by ultrafiltration (Amicon ultra 30,000 NMWL, Merck Millipore, Germany) with a buffer (20 mM HEPES-NaOH, pH 7.0, containing 100 mM NaCl and 0.05 % DDM) to remove retinal oxime and unreacted HA. To reconstitute the apo-proteins with ATR, the apo-proteins (∼ 1.7 μM) were incubated with 1.5 molar equiv. of ATR at 25 °C in the dark. The absorption changes were monitored during the retinal reconstitution.

### Structural modeling of SacRs

Structure of SacR_NC335_ was modeled by AlphaFold2 [11] using an unmodified version of ColabFold [12], with the default MSA pipeline, a MMseqs2 [31] search of UniRef [32] and environmental sample sequence databases [33, 34].

## Supplementary Figures

**Figure S1.**
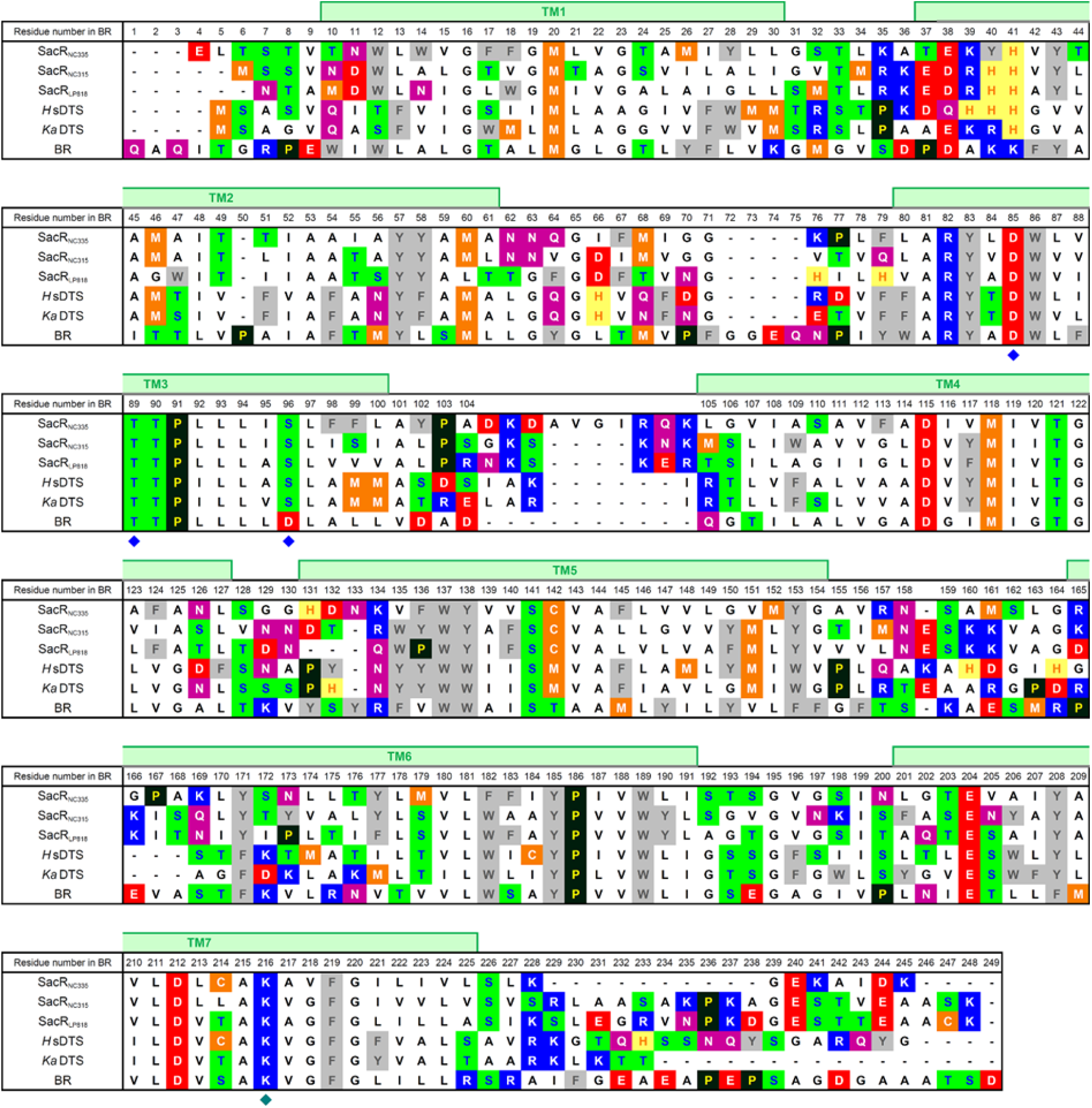
Amino acid sequence alignment of SacR with *H*sDTS, *Ka*DTS, and BR. The positions of the seven transmembrane helices (TM1–TM7) in the X-ray crystallographic structure of BR (PDB ID: 1M0L) are indicated by green rectangles. The triplet motif in TM3 and retinal-binding lysine are indicated by blue and green diamonds, respectively.

**Figure S2.**
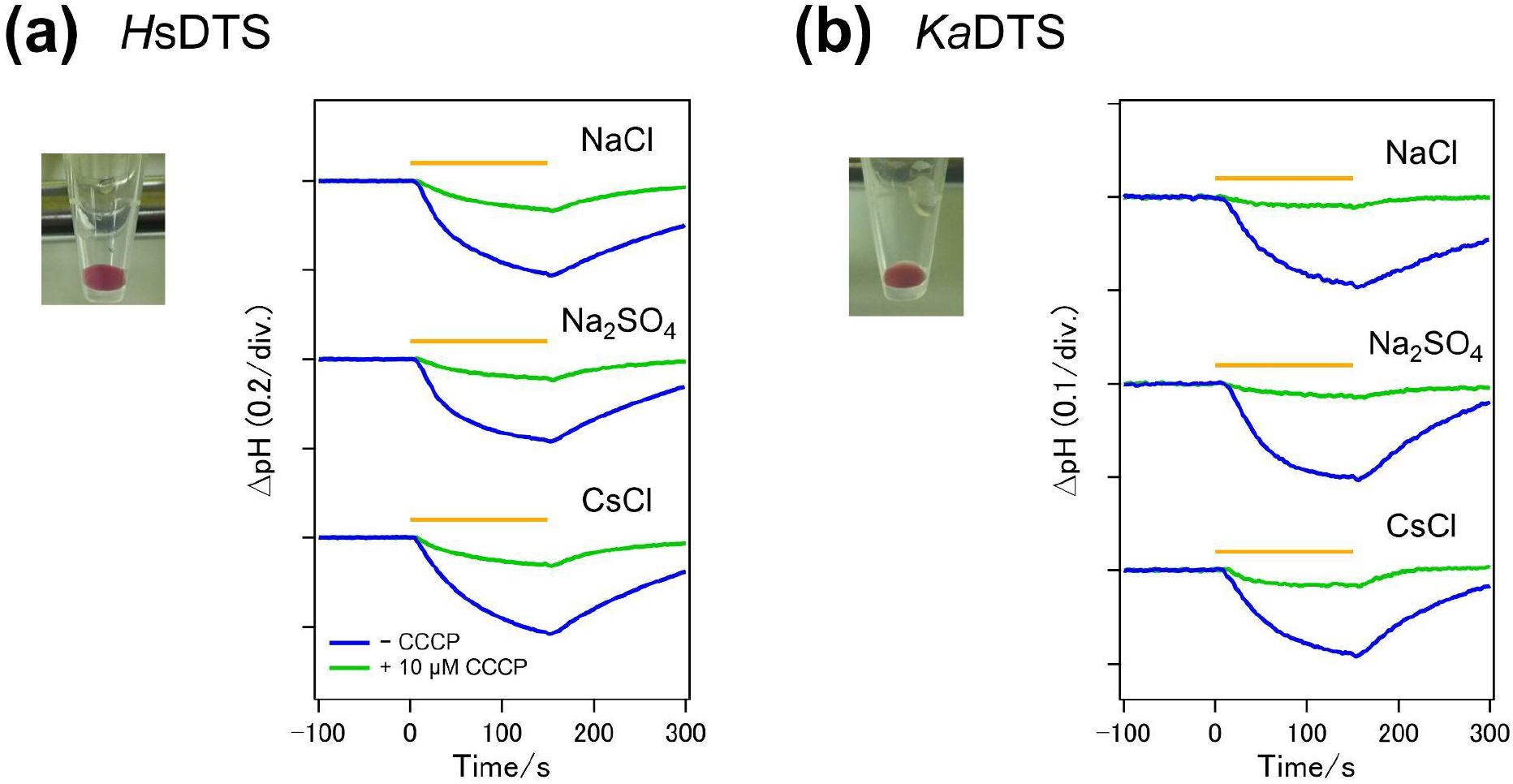
Proton transport assay of *H*sDTS (*Halomonas* sp.) and *Ka*DTS (*Kusheria aurantia*). Light-induced pH change in external solvent of *E. coli* cells expressing *H*sDTS (**a**) and *Ka*DTS (**b**) without (blue) and with (green) 10 μM CCCP. Light (*λ* ≥ 500 nm) was illuminated for a period indicated by yellow bars. Pictures of *E. coli* cell pellets expressing rhodopsins are shown.

**Figure S3.**
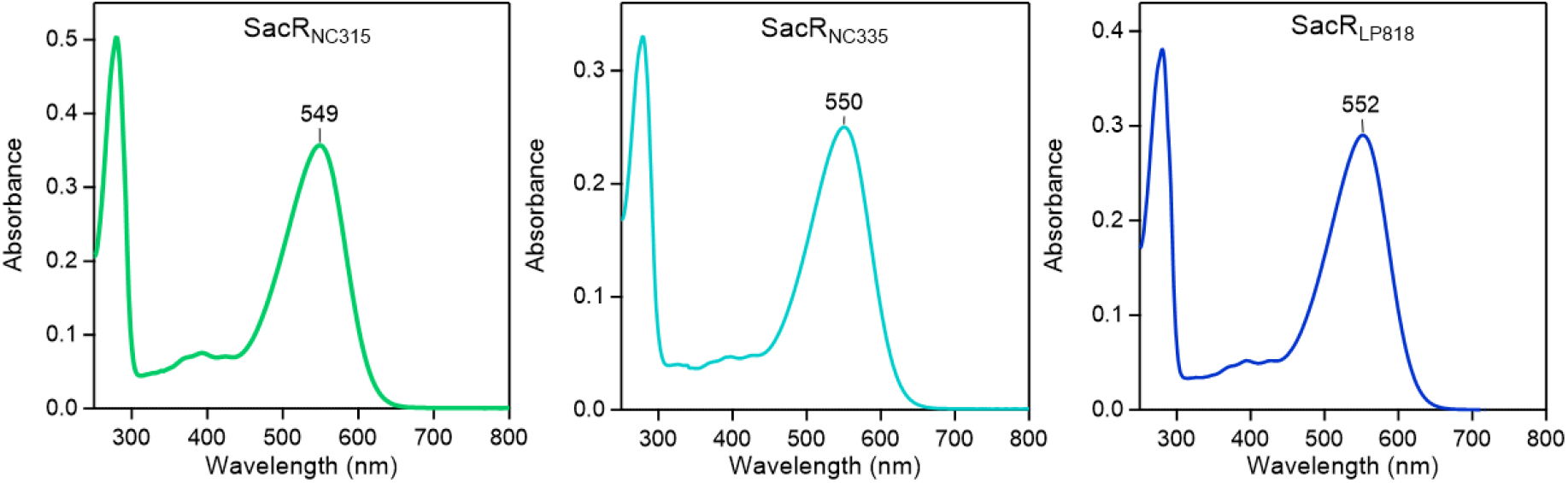
UV visible absorption spectra of SacRs in 100 mM NaCl, 20 mM HEPES–NaOH, pH 7.0, 0.05% DDM.

**Figure S4.**
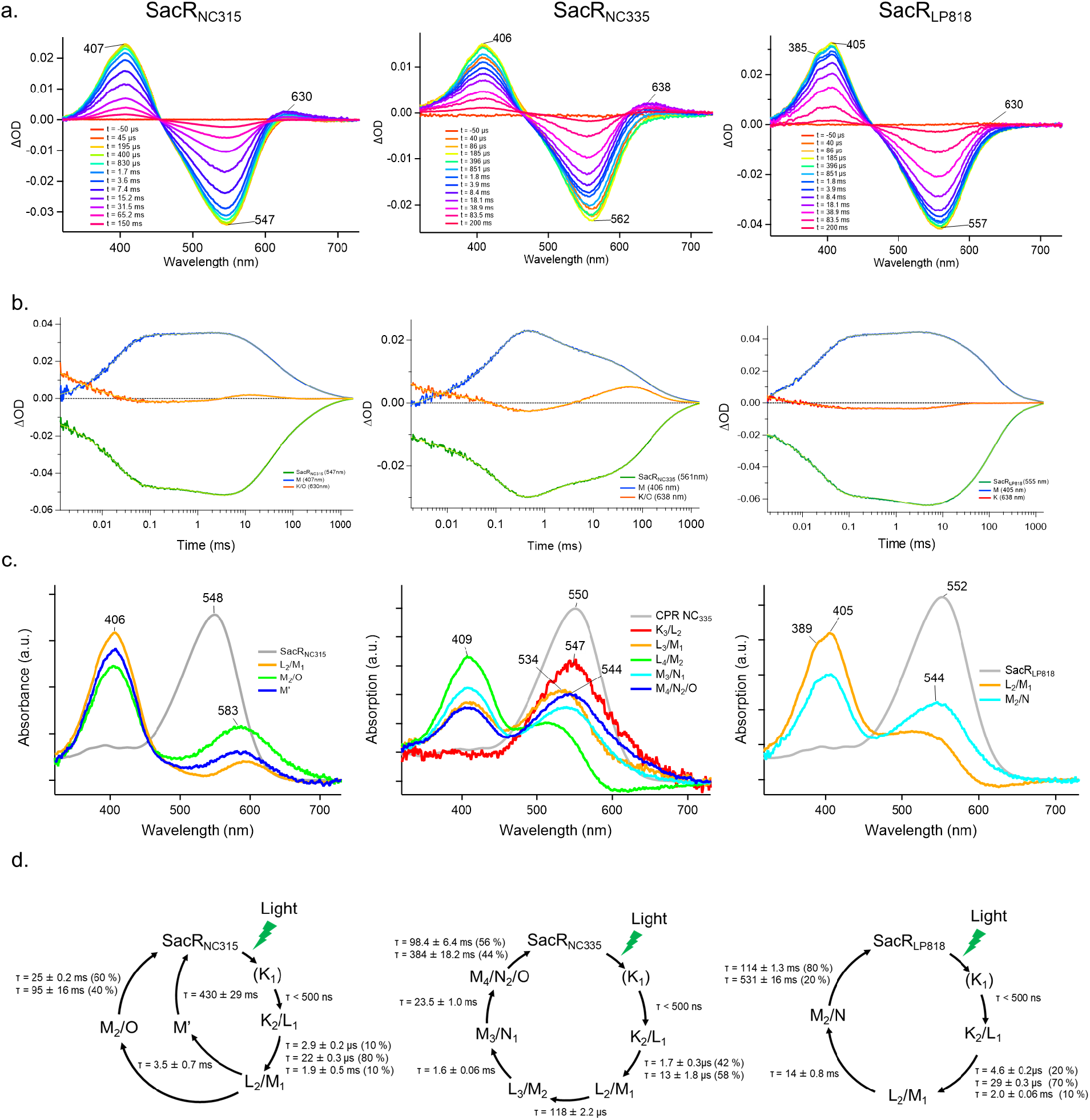
**a**) Transient absorption spectra of SacR_NC315_ (left), SacR_NC335_ (middle) and SacR_LP818_ (right). **b**) Time evolution of transient absorption change of SacR_NC315_ (left), SacR_NC335_ (middle) and SacR_LP818_ (right). The blue, green and orange lines represent the M intermediate, the breaching of the initial state, and the K or O intermediate, respectively. The yellow lines indicate fitting curves by multi-exponential function. **c)** Absorption spectra of photo-intermediates of SacR_NC315_ (left), SacR_NC335_ (middle) and SacR_LP818_ (right). The absorption spectra of photo-intermediates were calculated from the decay-associated-spectra obtained by multi-exponential fitting according to previous study [4]. **d)** Kinetic models assuming sequential photocycles of SacR_NC315_ (left), SacR_NC335_ (middle) and SacR_LP818_ (right) based on the fitting in **b**. The lifetime (*τ*) of each intermediate is indicated by numbers (mean ± S.D., the fraction of the intermediate decayed with each lifetime in its double exponential decay was indicated in parentheses).

**Figure S5.**
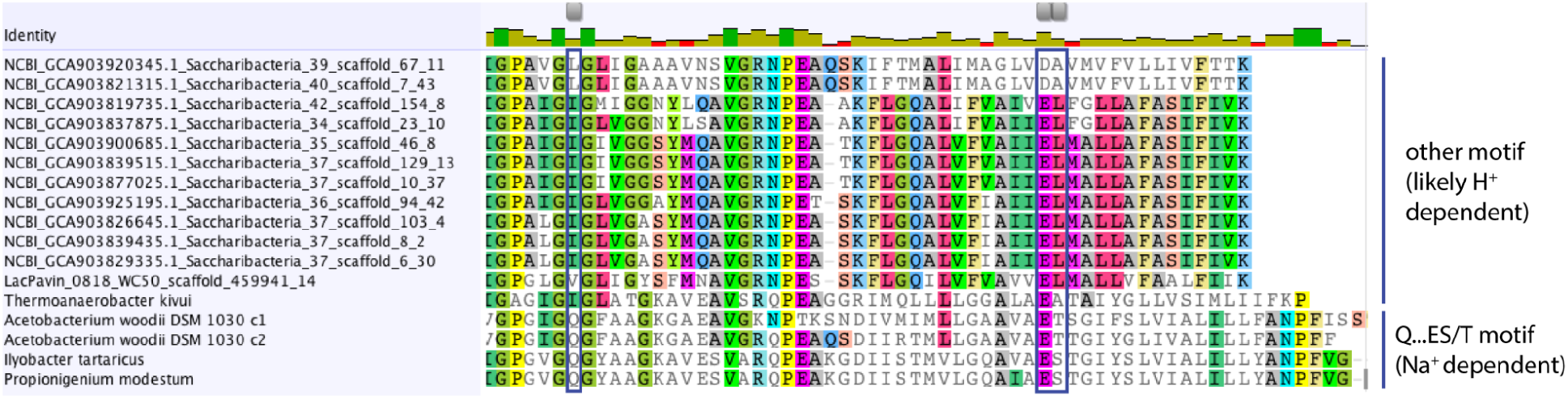
Partial protein alignment of the c subunit F_1_F_o_ ATP synthase from Saccharibacteria with rhodopsin and known references. Columns encoding the ion binding motif are indicated by blue boxes.

**Figure S6.**
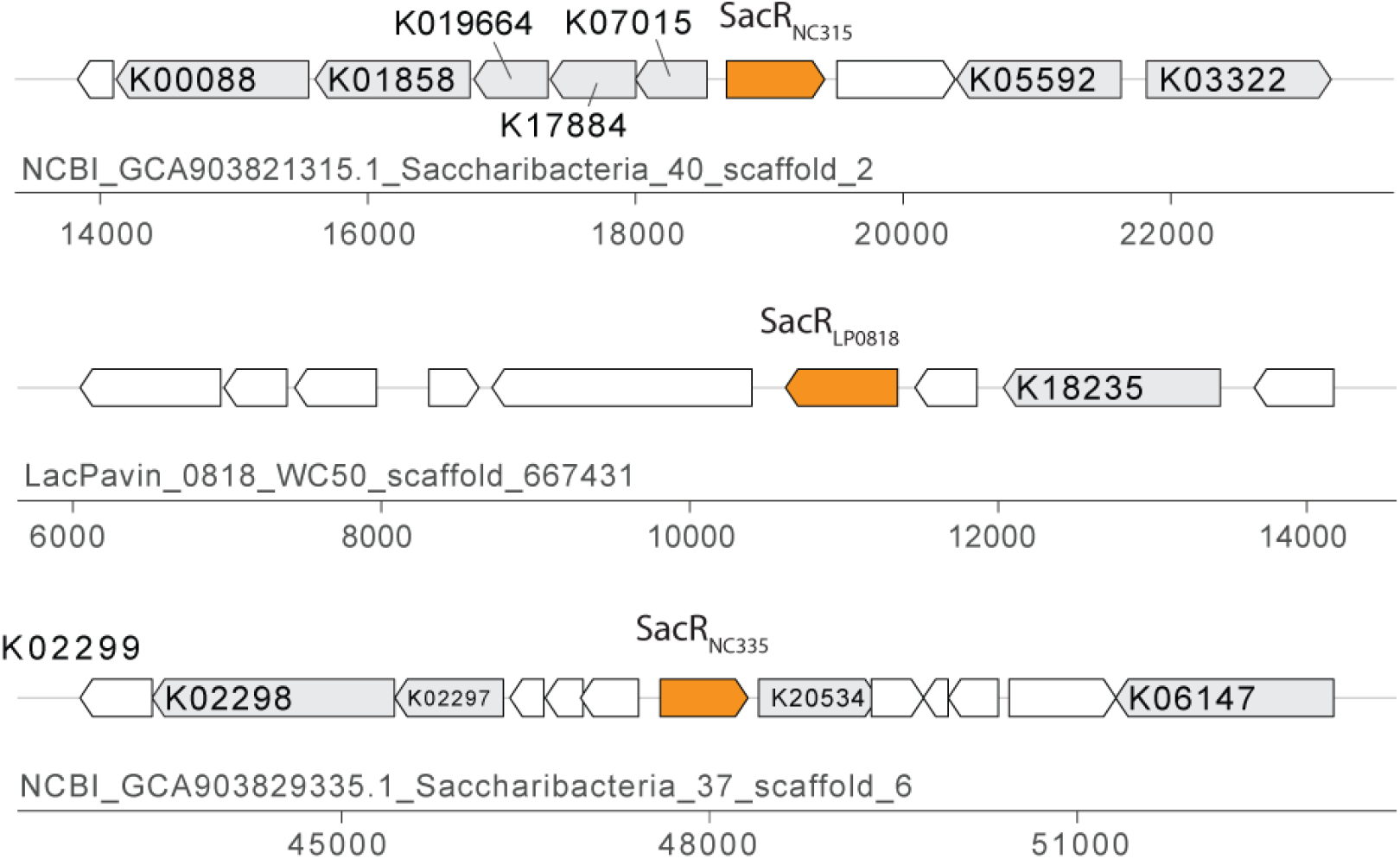
Immediate genomic context of the three characterized SacRs (orange). Neighboring genes with above-threshold KEGG annotations are indicated in gray with the highest-scoring HMM model. Genes without KEGG annotations are indicated in white.

**Figure S7.**
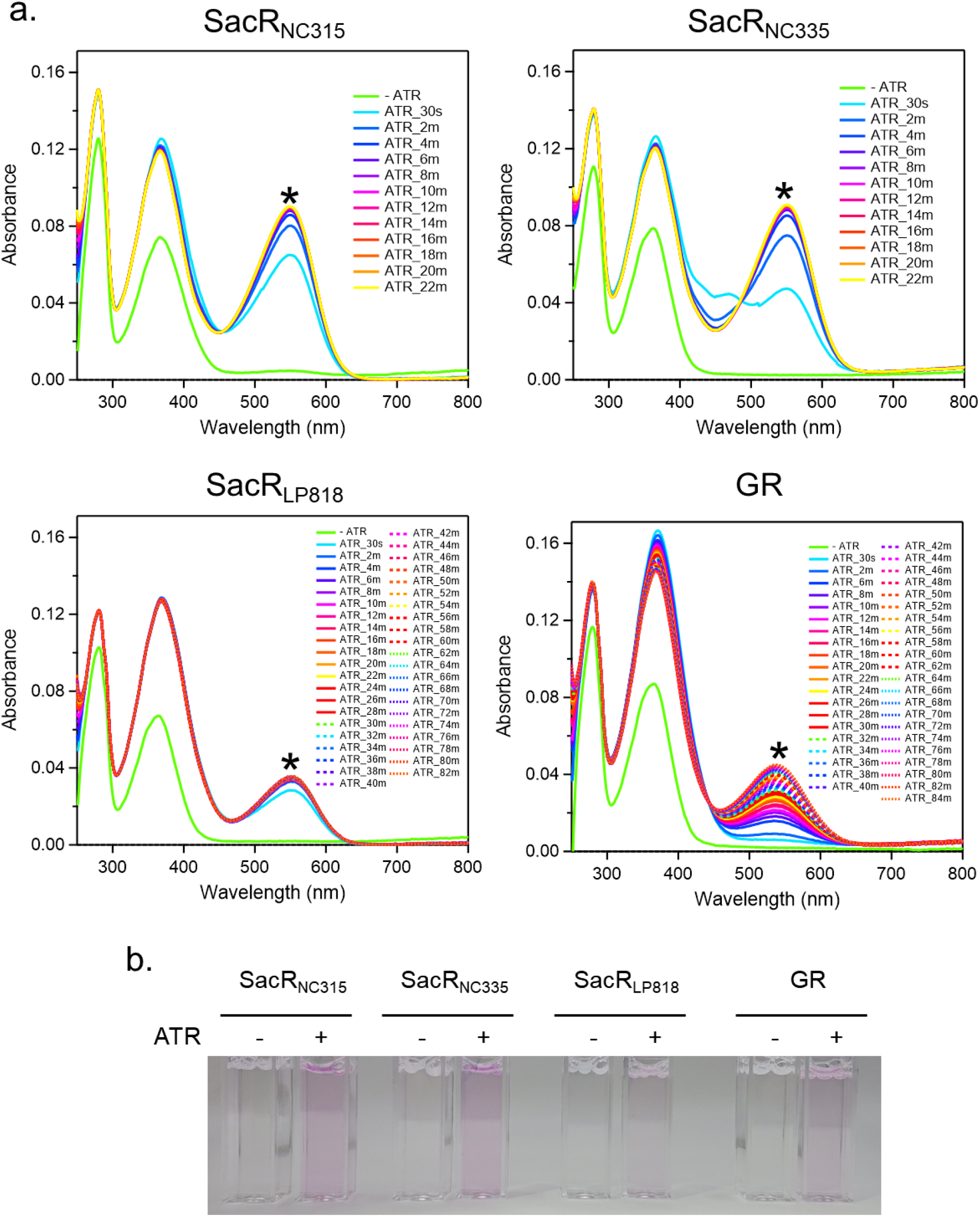
**a)** UV-visible absorption spectra showing the regeneration of retinal binding to SacRs and GR in 20 mM HEPES–NaOH, pH 7.0, 100 mM NaCl and 0.05% n-dodecyl-β-maltoside (DDM). The asterisks indicate absorption peaks of reconstituted rhodopsins. **b)** Appearance of SacRs and GR in solution without (−) and with (+) retinal (ATR). When supplemented with retinal, SacR and GR solutions were colored, indicating retinal regeneration.

**Figure S8.**
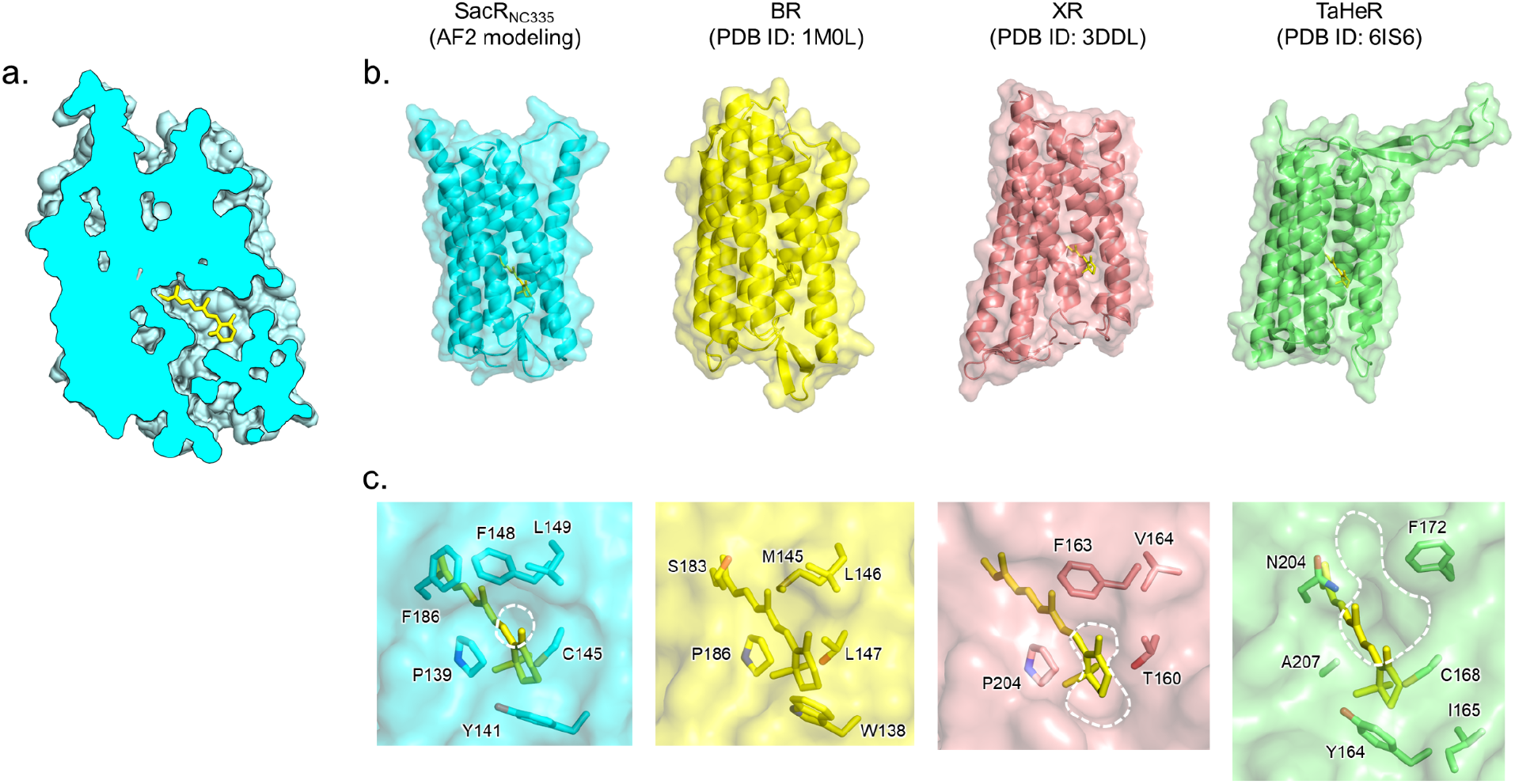
Modeled structure of SacR_NC335_. **a)** cross-section view of a SacR_NC335_ protomer modeled by Alphafold2 [11, 12]. **b**) Structural comparison of overall structures with other representative microbial rhodopsins. **c)** Magnified views of retinal-binding pockets with their molecular surfaces. The residues forming the β-ionone ring side of the retinal-binding pocket are indicated by stick models. White dashed circles represent the hydrophobic holes on the β-ionone ring side. No hydrophobic hole is present in the structure of BR.

**Figure S9.**
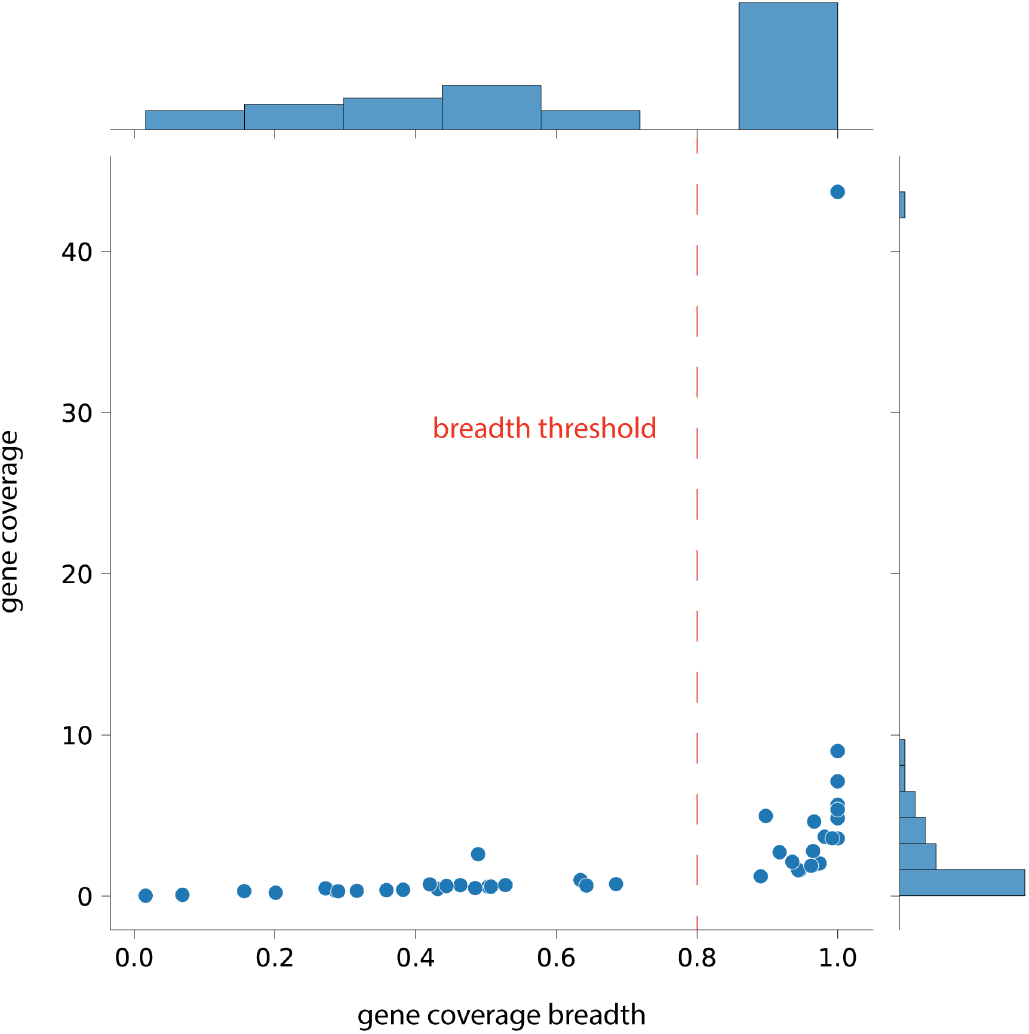
Sequencing read coverage and coverage breadth (fraction of gene covered by reads) for SacRs in analyzed freshwater metagenomes. Metagenomic samples in which any SacR obtained 0.8 coverage depth or greater were analyzed for genetic potential for retinal synthesis.

